# The Organizational Principles of Membranes Below 100 nm: Experimental Breakthroughs Occasion a “Modeling Manifesto”

**DOI:** 10.1101/292383

**Authors:** E. Lyman C.-L. Hsieh, C. Eggeling

## Abstract

New experimental techniques reveal the plasma membrane to be heterogeneous and "scale-rich," from nanometers to microns, and from microseconds to seconds. This is critical information, as heterogeneous, scale-dependent transport governs the molecular encounters that underlie cellular signaling. The data are rich, and reaffirm the importance of the cortical cytoskeleton, protein aggregates, and lipidomic complexity to the statistics of molecular encounters. Moreover, the data demand simulation approaches with a particular set of features, hence the “manifesto”. Together with the experimental data, simulations which satisfy these requirements hold the promise of a deeper understanding of membrane spatiotemporal organization. Several experimental breakthroughs are reviewed, the constraints that they place on simulations are discussed, and the status of simulation approaches which aim to meet them are detailed.

## Introduction

The organization of the plasma membrane, and functional consequences thereof have been discussed since at least the 1970s. Motivated by the phase behavior of cholesterol/phospholipid mixtures,(1) Jain and White suggested the possibility of “lipid islands” — solid landmasses anchored in a sea of lipids.(2) This was a strictly structural model, no functional role was attributed to these islands. Twenty years later, Simons and Ikonen introduced “lipid rafts”: protein was added to lipid islands, the assembly was set adrift, and functional consequences were attributed, by virtue of the raft’s ability to regulate spatiotemporal encounters.(3) Then followed substantial effort to biophysically characterize ternary (and sometimes more elaborate) lipid mixtures.(4) During this period, the equilibrium phases that are observed in such mixtures were hypothesized to provide a thermodynamic basis for rafts in live cells.

The raft concept has evolved into a much more nuanced view,(5, 6) but is not universally accepted, for at least two reasons. First, because direct imaging of potential rafts in living systems with nanometer spatial resolution is challenging, a truly decisive experiment (for or against rafts) has not yet been demonstrated. Second, the raft idea has spawned very few precise, quantitative predictions for *functional consequences* that are amenable to experimental test. We view the latter point not as a failure of the raft model, but a failure of model*ing*, or more precisely, a failure of modeling to intersect usefully with experimental measurement.

This Perspective therefore addresses the intersection of modeling and experimental measurement in the context of plasma membrane organization. Our aim is not to review all work on membrane heterogeneity, and we will consequently only cite exemplary work (and apologize to those many others that we fail to mention). Still, this review is occasioned by significant advances in experimental measurement, which in turn place particular demands on modeling approaches, and offer a new opportunity for a dialogue between modeling and experiment. Recent, transformative experimental advances for studying cellular membrane organization and dynamics include:

i. Single particle tracking (SPT) especially with increased sensitivity due to combination with photoswitching (7) or contrast enhancement using scattering(8) or coherent illumination and interference (such as interferometric scattering (iSCAT) (9-11) and coherent brightfield (COBRI) imaging microscopy(12) to image lipid and protein dynamics within and between membrane compartments.
ii. Fluorescence correlation spectroscopy(13) (FCS) for molecular dynamics studies, especially when combined with spot-variation,(14) camera-based detection,(15) image correlation approaches (16), near-field microscopy(17) or super-resolution STED microscopy (STED-FCS) (18).
iii. Super-resolved imaging based on electron(19), fluorescence (such as STED or (f)PALM/(d)STORM microscopy)(5) or near-field microscopy(20) to map molecular membrane organization in living and fixed cells with improved spatial resolution.
iv. Environment-sensitive fluorescent dyes (such as C-Laurdan), reporting for example on the membrane polarity or lipid packing (e.g. (21)).
v. Model membrane systems such as supported and black-lipid bilayers or giant-unilamellar vesicles (GUVs), whose composition can be specifically tuned, as well as cell-derived giant-plasma-membrane vesicles (GPMVs), which contain the lipid and protein diversity of the original cell. Especially valuable are phase-separated forms thereof, which enable indicating influences of lipid membrane order on membrane organization(22).
vi. Improvements in general structural and biochemical analysis tools, e.g. based on mass spectroscopy/lipidomics, NMR or related techniques(6).

We begin from an agnostic view, and list a few observations about plasma membrane organization that are not likely to be controversial. This will set the stage to explain the significance of the aforementioned experimental developments, which will in turn identify features with which a modeling approach must contend.

### The lipid fraction of the plasma membrane is a diverse mixture of hundreds of different lipids

Measurement of the chemical composition of lipid samples is now possible with exquisite sensitivity, achieved by electrospray ionization and tandem mass spectrometry.(23–25) These measurements reveal a typical plasma membrane to be comprised of hundreds of distinct lipid species, differing in every aspect of their chemistry. Without ascribing any functional significance, such a mixture is *expected* to be heterogeneous in composition — it is extremely unlikely that a mixture of 100 different components will constitute an ideal mixture.

In animal cell membranes, 20-40% by mol of the total lipid is cholesterol. In lipid mixtures much simpler than the plasma membrane, cholesterol tends to order hydrocarbon chains and increase area density of lipid, while also fluidizing already highly ordered saturated-lipid environments. At fixed composition, a miscibility transition is observed; and at temperatures typically below physiological temperature, coexisting liquid-ordered (L_o_) and liquid-disordered (L_d_) phases are observed. These statements are true of mixtures derived from plasma membrane as well (such as in GPMVs), although the ordering/packing effect of cholesterol seems to be weaker, the two phases are more similar to one another than in ternary mixtures, and the miscibility temperature varies over tens of degrees.(26)

### Between 15 and 30 % of the cross sectional area of the plasma membrane consists of integral membrane proteins

The precise estimate varies depending on the methodology, but it is clear that a significant fraction of the membrane is occupied by protein. Furthermore, the distribution of membrane protein is not uniform, with many types of membrane protein observed to form clusters and aggregates. (5, 6)The mobility of membrane proteins is also not uniform, as indicated by fluorescence based SPT.(8, 27)

### The cortical cytoskeleton partitions the membrane

The SPT measurements of Kusumi and coworkers showed that in live cells cortical actin acts as a partial barrier to membrane protein diffusion on length-scales of tens to hundreds of nm. (8, 28) This “picket fence” model is confirmed by more recent STED-FCS measurements(29) and correlative SPT and (super-resolved) imaging experiments(30). In model membranes, modeling studies and comparison to imaging experiments such as with STED microscopy revealed coupling between lipid phases and reconstituted cortical actin networks.(31–33) Though much of the literature focuses on cortical actin, other structural proteins may have a role in membrane partitioning, such as spectrin and septins.

It is clear from the above discussion that the membrane is “scale-rich,” and heterogeneous over a wide range of length and timescales, due to very different mechanisms — lipid mixing at short length-scales, protein aggregation at intermediate length-scales, and the picket fence mechanism at longer length-scales. *Measurements of lipid and protein motion in membranes are therefore length-scale and timescale dependent* — the motion observed on 20 nm length-scales need not be the same as that observed on 100 nm length-scales. This is why the experimental techniques mentioned above represent critical advances, demanding a new look at modeling approaches. In the remainder, we discuss in more detail some selected recent experimental measurements, mainly those based on SPT, STED-FCS and iSCAT/COBRI, and then turn attention to modeling approaches.

### Brief review of experimental techniques

#### High speed single particle tracking

SPT has yielded tremendous insights into membrane organization by allowing the direct observation of hindrances in molecular diffusion maps. As highlighted before, the Kusumi lab used video microscopy of membrane components with nm spatial resolution at 40,000 Hz to show that the membrane is compartmentalized by the cortical actin cytoskeleton. This transformational advance showed unequivocally that the membrane is partitioned into compartments with a lengthscale on the order of tens to hundreds of nm, as revealed by the hop diffusion of individual lipids tagged by 40 nm gold nanoparticles, with the size of compartments varying across different cell types.(8) Apart from fundamental insight into the organization of the plasma membrane, these measurements demonstrated the need for modeling alongside measurement, in order to distinguish compartmentalization from statistical fluctuations intrinsic to stochastic trajectories.

#### FCS measurements of lipid diffusion in membranes

By determining average transit times of molecules through the microscope’s observation spot, FCS allows the determination of molecular mobility in model and cell membranes(13). Recording FCS curves at different spot sizes (so-called spot-variation (sv) FCS) allows different diffusion modes to be distinguished, such as free diffusion, trapped diffusion due to transient halts at molecular complexes or domains, and hop diffusion(14). Yet, due to the limited spatial resolution of confocal microscopy, conventional svFCS recordings can only interpret the diffusion characteristics at the relevant molecular scales by extrapolation of the acquired results to the nanoscale(14). A remedy is to use optical microscopy approaches with enhanced spatial resolution or reduced observation spot sizes. Subdiffraction excitation volumes can be realized by near-field approaches, where the sample (such as a cell) is placed very close (within nanometer distance) to nanometer large, excitation-light guiding apertures. Near-field FCS data has confirmed and further detailed diffusion modes in model and cellular membranes as discovered by the aforementioned svFCS experiments(17, 34, 35), yet the close proximity of microscope equipment may bias molecular diffusion characteristics, prevent measurements inside cells or further away (e.g. apical) membranes, and makes spot-variation measurements more tedious, since it requires separate experiments with different apertures. A remedy are super-resolution optical microscopy approaches such as STED microscopy. As in conventional microscopes, STED microscopy employs lenses for the focusing of the fluorescence excitation laser light, and the sample can be placed micrometers away from any optical element (into the optical far-field). Modulation of the fluorescence emission in the focused laser spot now allows creating efficient fluorescence observation spots of sub-diffraction (<<200 nm) sizes. Consequently, FCS can now be employed with observation spots that may in principle be reduced without limit (practically, due to signal-to-noise issues, down to around 30 nm in diameter). In addition, the observation spot can straightforwardly be tuned through changing laser power levels, allowing access to sub-diffraction svFCS experiments(18, 36, 37), even within a single measurement when using gated detection approaches(38). Using this STED-FCS approach, diffusion modes of various fluorescent lipid analogs and proteins in model membranes and the plasma membrane of living cells were observed with unprecedented spatiotemporal resolution(14,53,54). For example, it was shown that the diffusion of sphingolipids was hindered through transient halts (trapped diffusion) from interactions with immobilized or less mobile membrane entities, while such hindrance was not observed for glycero-phospholipids (14,53). Simultaneous acquisition of STED-FCS data at different spatial positions by repetitively scanning a circle or line (scanning STED-FCS, sSTED-FCS) highlighted a non-continuous distribution of such interaction sites; these hot spots were located at distinct locations, on average 150-200 nm apart, <80 nm in size, and transient on the second time scale(39).

What may be the nature of the inhomogeneities observed by STED-FCS in cells? Detailed STED-FCS studies using drug treatments or on GPMVs disclosed a dependency of the trapped diffusion on cholesterol and the cortical actin cytoskeleton, indicating that the (more immobile) binding partners are linked to the actin cortex and the interactions are mediated by cholesterol(40, 41). Interestingly, the trapped diffusion of ganglioside, in contrast to that of sphingolipids, was much less, if at all dependent on cholesterol and the actin cytoskeleton, probably due to their large polar head-groups(40, 41). Experiments on model systems may provide further clues. For example, STED-FCS data in phase-separated ternary membrane ^bilayers has indicated an absence of hindered diffusion, both in the Lo and Ld phases,^ at least down to the probed 40-nm length scales.(42) Coupling of the model membranes to a cortical actin cytoskeleton, however, introduced hindrances in diffusion, mainly due to the aforementioned coupling between lipid phases and the reconstituted cortical actin networks(32, 33). In contrast, no correlation has been observed between Lo-partitioning characteristics of lipid analogs and their diffusion modes in living cells, questioning the viability of the direct transfer of results obtained on model membranes to the live-cell case(39).

#### iSCAT and COBRI based measurements of spatiotemporal dynamics

iSCAT enables detection of weakly contrasting objects by interfering a coherent reference beam with the weak signal of interest.(43, 44) de Wit, *et al.* have reported label-free, direct imaging of freely diffusing solid-ordered domains at a droplet interface bilayer.(11) This is a type of supported bilayer geometry, in which a bilayer is formed at the interface of two monolayers, one of which sits on a hydrogel atop a support. Using this setup, de Wit, *et al.* could directly detect 200 nm large nanodomains and track their motion with a localization precision of 10 nm. The motion of smaller domains is also observable as variation in contrast. Notably, their size is estimated based on a calibration obtained for larger domains and extrapolated based on a model for the size dependence of domain diffusion. (This is revisited below.)

iSCAT also offers a unique approach to label-based SPT of individual lipids and proteins. At a laser intensity of 100 μW/μm^2^, the location of a lipid chemically linked to a small (20 nm dia.) gold nanoparticle may be detected with a spatial resolution of 3 nm and a time resolution of 20 μsec (Fig. 2A).(9, 10) Since there is no photobleaching, very long trajectories of individual particles may be observed — tens of thousands of steps, with total durations of a few seconds. One of us (CLH) has recently demonstrated that an individual lipid (biotin-cap-DPPE labeled by 20 nm gold nanoparticle) switches from normal diffusion to subdiffusion when crossing a boundary from an L_d_ phase to an L_o_ phase, using a 40:40:20 mixture of DPPC/diPhy/Chol on a mica support (Fig. 2A).(9) Trajectories in the L_o_ phase show subdiffusion up to approximately one msec, after which the diffusion becomes Fickian. Statistical analysis of particle trajectories suggests transient confinement, perhaps in nanoscopic substructures previously observed in molecular dynamics simulations in similar lipid mixtures.(45, 46) Very recently, similar nanoscopic substructures in L_o_ phase and its influence to lipid diffusion were also detected experimentally by FCS with specially designed planar plasmonic antennas.(35)

**Figure 1.**
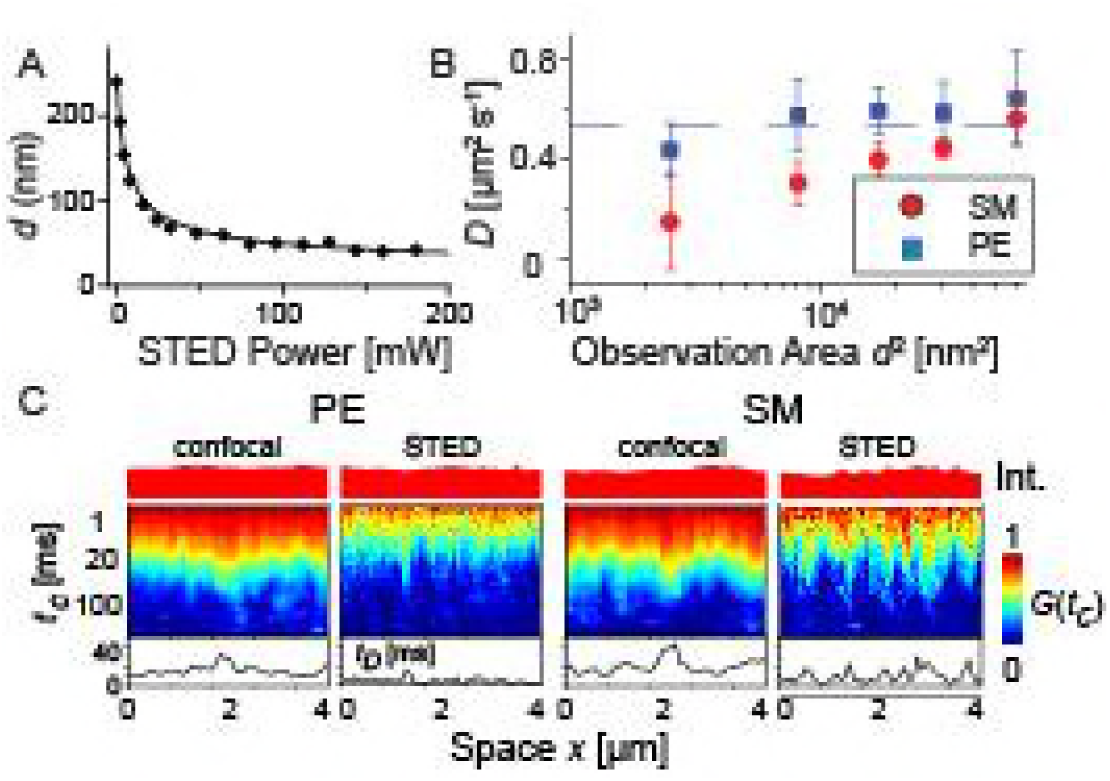
Length-scale-dependent diffusion characteristics of fluorescent lipid analogs observed by STED-FCS in sub-diffraction sized observation spots. (A) STED-FCS exploits the phenomenon that the size of the observation spot (diameter d) can be tuned at sub-diffraction scales by the power of an additional STED laser (adapted from ref (37). Super-resolution optical microscopy of lipid plasma membrane dynamics. Essays Biochemistry. 57:69–80). (B) Dependence of the apparent diffusion coefficient D as determined by STED-FCS on the observation spot sizes (diameter d squared) for a fluorescent phospholipid (PE, blue) and sphingomyelin (SM, red) analog in the plasma membrane of live PtK2 cells, indicating close to free diffusion (dashed line) for PE and hindered trapped diffusion for SM. (C) sSTED-FCS data of the same measurements on PE and SM: Correlation carpets of PE (left two panels) and SM (right two panels) diffusion of confocal and STED recordings (as labeled) for 10-s measurement time: Total intensity (respective upper panels, normalized), correlation carpets (respective middle panels) and fitted transit times t_D_ (respective lower panels) over space, i.e. for the pixel positions of the scanned line (x-axis). The correlation carpets show the normalized temporal correlation curves corresponding to the correlation amplitude G(t_c_) depicted by the color code, while the correlation time t_c_ is encoded in the y-axis. The respective transit times t_D_ for every pixel on the scan trajectory can be estimated from the color decay (yellow regions). Large fluctuations in transit times (t_D_) indicating positions of hindered diffusion are only revealed for SM in the STED recordings. The sites of hindered diffusion were smaller than 80 nm and transient, i.e. they changed over consecutive time windows. (Adapted from ref 39).

**Figure 2.**
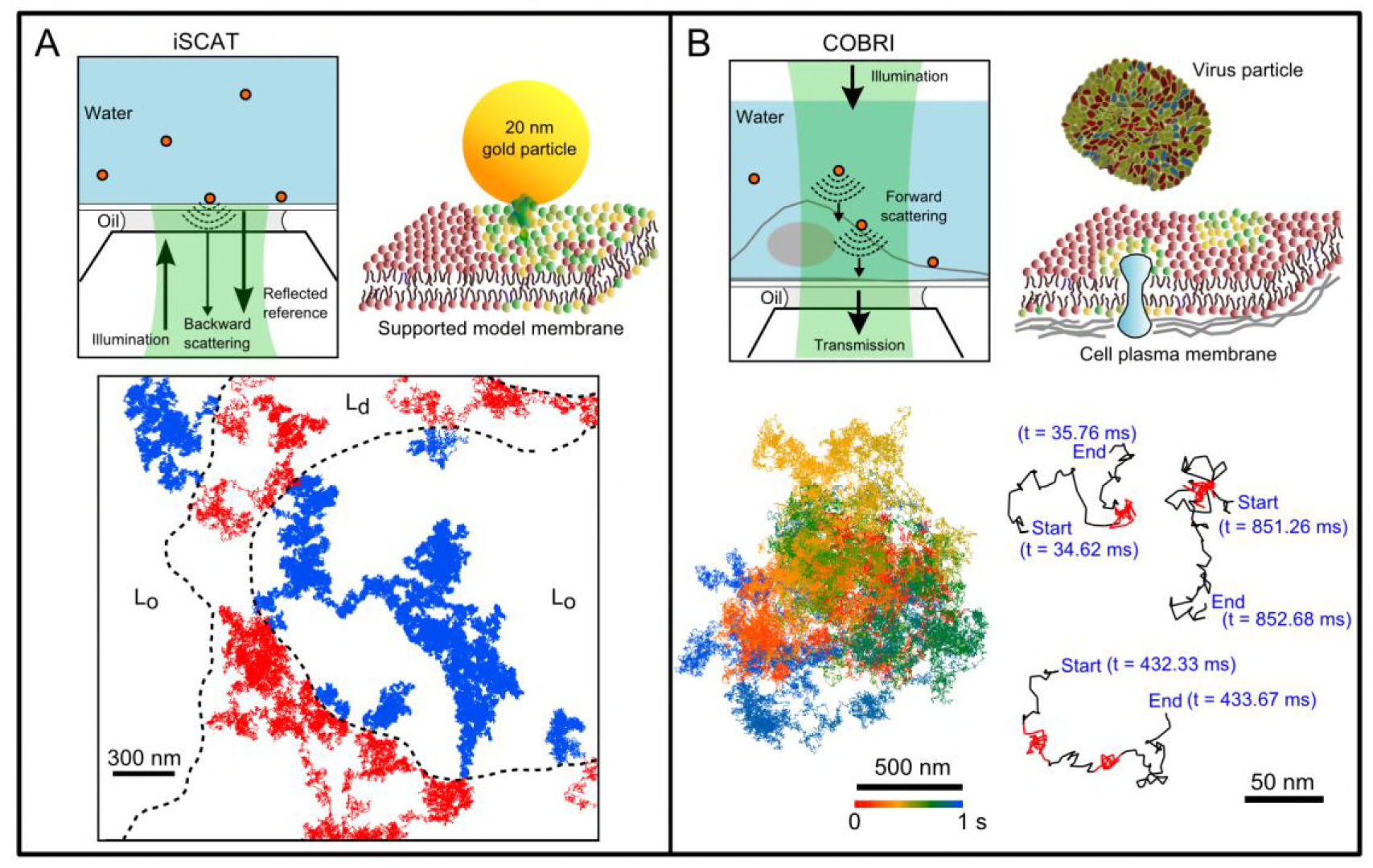
iSCAT and COBRI microscopy enable high speed imaging with nm spatial localization precision. (A) In iSCAT, coherent illumination and detection from below the sample observes a lipid tagged with a 20 nm gold particle with nm spatial localization precision and 20 μsec time resolution. The lipid switches from normal diffusion in the liquid-disordered phase (red portion of trajectory) to subdiffusion in the liquid-ordered phase (blue trajectory). (B) COBRI is the transmission counterpart of iSCAT, which typically operates at much lower light intensity at the price of reduced sensitivity. The trajectory (bottom right) shows a virus particle diffusing on the surface of a HeLa cell with a 100,000 Hz frame rate, revealing very fast surface diffusion and transient localizations immediately after landing. Adapted with permission from (9) and (12).

Recently, Hsieh and coworkers demonstrated “COBRI” microscopy, the transmission counterpart of iSCAT (Fig. 2B).(12) Both iSCAT and COBRI detect linear scattering light (backward scattering in iSCAT and forward scattering in COBRI) from nano-sized objects via interference. The signal-to-noise ratio (SNR) of iSCAT and COBRI is determined by the number of detected signal photons, independent of the electronic noise, giving optimal sensitivity in high speed measurements.(47) Using COBRI and proper image processing, single 40 nm gold nanoparticle were tracked with 3-5 nm precision at high speed.(48) The transmission geometry of COBRI makes it less sensitive to membrane fluctuation and thus easier to observe small particles in living cells. Using COBRI, Hsieh and coworkers reported high speed tracking (100,000/sec frame rate) and 3D localization (1.7 – 2.3 nm precision) of individual vaccinia virus particles as they land and then diffuse laterally on the surface of a live cell, with total acquisition times of seconds (Fig. 2)(12). The measurements clearly show very heterogeneous diffusion, with virus particles sampling zones of transient confinement on submillisecond timescales, eventually (within seconds after landing) becoming strongly confined. By obtaining trajectories tens of thousands of steps long, the coherent illumination detection methods alleviate some of the statistical limitations inherent to SPT approaches, enabling measurements uniquely suited to the scale-rich environment of cell membranes.

#### Challenges

For both iSCAT/COBRI and STED-FCS, it is necessary to specifically label molecules of interest, which is laborious and may influence the motion of interest, especially when using large gold-nanoparticle tags as in most iSCAT/COBRI experiments.(49) Recent development of sensitivity enhancement in iSCAT demonstrates the capability of seeing single biological macromolecules directly in aqueous environment without any labels, which shows the promise of revealing single membrane protein dynamics in their most native form(50, 51). The STED-FCS approach admits the use of much smaller labels, and is ideally suited to lengthscale-dependent measurements of diffusion, but being a rather local measurement (even for the lines in the sSTED-FCS recordings), does not easily provide information on heterogeneity across the cell surface. However, improvements in image-based correlation approaches may provide a remedy.(16, 52) In addition to the possible effects of labeling, for both iSCAT/COBRI and STED-FCS additional laser illumination and/or stronger laser light levels are usually implemented for measurements at higher spatiotemporal resolutions (high laser intensities inherently required due to limited optical cross-sections of the available labels, such as scattering probes in iSCAT/COBRI, and fluorescent labels in STED-FCS). Unfortunately, intense illumination may potentially induce photo-toxic effects (such as cell death), which bias the observations and need to be carefully considered. Therefore, control experiments are required, e.g. making use of different observation techniques with different label types and illumination levels, or of complementary samples such as model membrane systems. As different techniques have their advantages and limitations, it is always valuable to probe the same phenomena with multiple techniques. Nevertheless, recent advances in recording schemes (such as for STED microscopy imaging)(53) are guiding ways for reducing phototoxic effects.

### Modeling Requirements: A Manifesto

Most of the measurements described above observe the motion of a protein or lipid, and then infer underlying membrane structure and interactions from changes or differences in mobility. Conversely, if a simulation approach is to obtain predictive validity regarding the encounter of membrane proteins and/or lipids, it must correctly model *mobilities*, and how they depend on the complications — such as a heterogeneous distribution of protein and lipid, coupling to the cytoskeleton — described in the introduction. It is therefore essential that a modeling approach reproduce quantitatively the mobility of lipids and membrane proteins across a variety of experimental systems, from simple to complex.

Even in a simple membrane — fluid and homogeneous, infinite in extent, and bounded above and below by an infinite bulk of solvent — the mobility of a membrane protein is nontrivial. Treating the membrane as an incompressible, low Reynolds number fluid bounded by bulk solvent of significantly lower viscosity, Saffman and Delbruck (SD) derived an expression for the mobility of a cylindrical membrane inclusion.(54) In the SD model, the ratio of the viscosity of the membrane to that of the surrounding fluid introduces a new lengthscale (referred to in the following as L_SD_(54)), which marks a crossover in the hydrodynamics, from 2D behavior governed by the membrane viscosity to 3D behavior governed by the viscosity of the bounding solvent. For homogeneous fluid phase membranes, L_SD_ ranges from 100 to 1000 nm. Comparison to experimental measurements shows that the SD model accurately captures the dynamics of membrane proteins.(55)

For the quasi-2D diffusion that is characteristic of proteins in membranes, qualitatively different results are obtained for “free-standing” membranes — that is, bounded above and below by an infinite extent of solvent — and other boundary conditions. Motivated by experiments by Falke and coworkers,(56) Camley and Brown showed that the logarithmic dependence of mobility on protein or domain radius that is predicted by SD is lost for membranes on a solid support, with mobility instead depending linearly on radius for oligomeric protein assemblies.(57) This result reflects the loss of symmetry across the membrane midplane — momentum is dissipated by the support, while on the other side it is free to propagate into the solvent. It is therefore a mistake to use SD theory (or any other theory or simulation based on a free-standing membrane) to interpret experimental results obtained for a supported bilayer — any agreement is fortuitous, and any contradiction is irrelevant. In this light, recent iSCAT measurements of domain diffusion(11) bear reconsideration, using the approach of Camley and Brown.(57) Observed protein mobility also depends strongly on the presence of an immobile fraction of lipids or proteins, which may result from interactions with the cortical cytoskeleton. Oppenheimer and Diamant have shown that these may be very significant effects even at very low area fraction for a random configuration of immobile species (Fig. 3B), with strikingly counterintuitive results, such as mobilities that are either enhanced or reduced depending on the direction of the motion relative to the immobile species.(58, 59) The immobile fraction in a plasma membrane may not satisfy either requirement.

**Figure 3.**
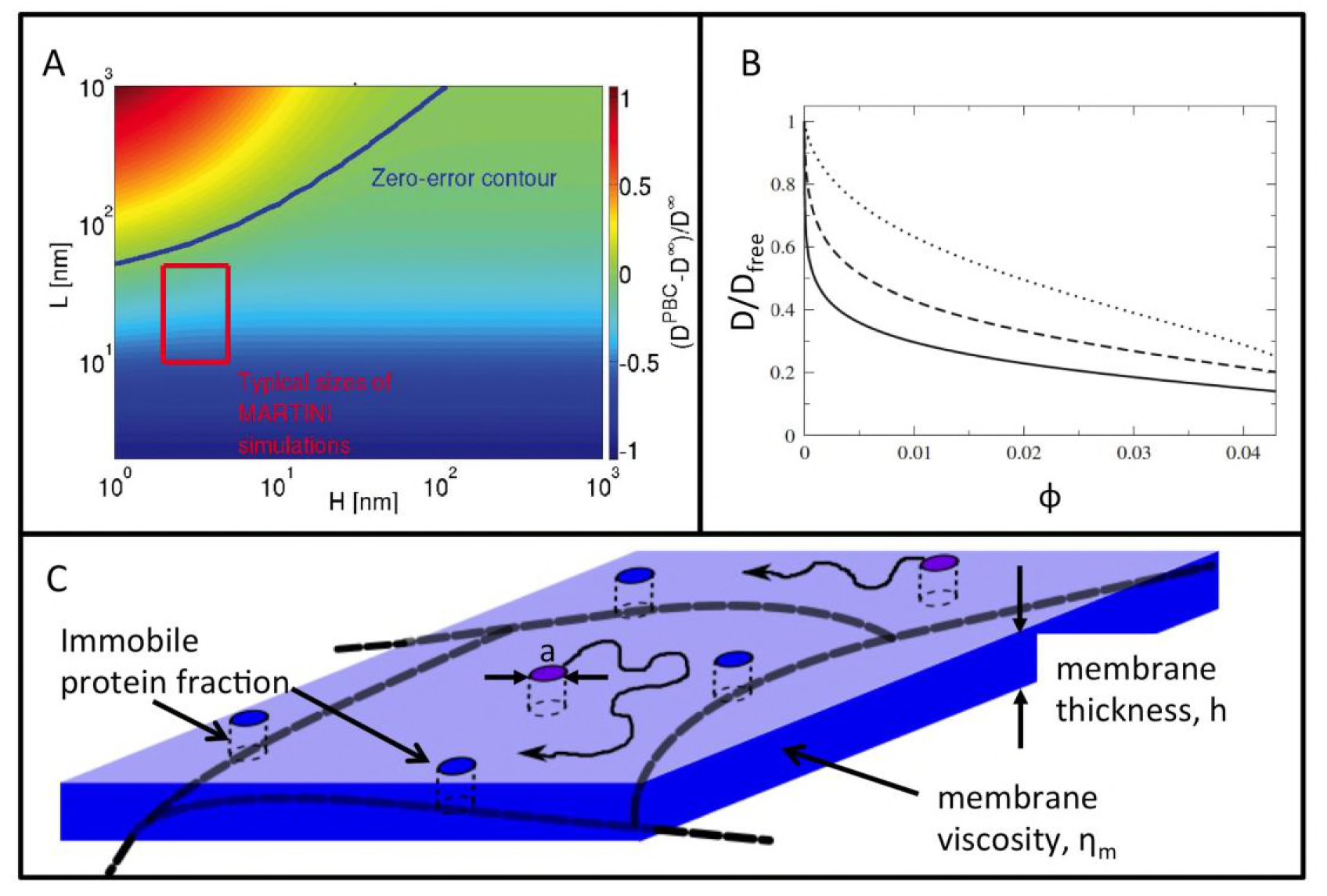
Membrane hydrodynamics are long-ranged, and often counterintuitive. (A) Extension of the SDHPW theory to systems with periodic boundary conditions reveal significant finite size artifacts for molecular dynamics simulations. The diffusion constant of an integral membrane protein in a simulation (D^PBC^) may be larger or smaller than that of the same protein in an infinite system (D^∞^), depending on the size of the system in the directions normal to the membrane (H) and in the plane (L). Adapted with permission from (59). (B) Even at very low area fraction ϕ, a randomly arranged set of immobile proteins strongly impacts the diffusion D of mobile proteins, relative to diffusion in the absence of an immobile fraction (Dfree), depending on the radius a of the mobile proteins. Shown are the dependence for a/L_SD_ = 10^−3^ (solid), 10^−2^ (dashed), and 10^−1^ (dotted). Reprinted with permission from (57). (C) The mobility of a mobile protein (pink cylinder) in a simulation or experimental measurement depends on a nonrandom assortment of immobile proteins (blue) that are bound to the cytoskeleton, as well as other complications such as local variations in membrane viscosity η_m_.

More recently, Camley et al. have obtained a generalization of the SD result for systems with periodic boundary conditions as commonly employed in molecular dynamics simulations (in the following called periodic Saffman-Delbruck or “PSD”), and verified the PSD theory by molecular simulations.(60) Because the dimensions typical of molecular dynamics simulations are of the same order as L_SD_, observed mobilities are contaminated by a significant finite size effect, resulting in simulated diffusion coefficients that may be larger or smaller than their infinite system counterparts by 100% or more (Fig 3A). This result has already been extensively confirmed by additional molecular dynamics simulation.(61, 62) If the goal of a simulation is to quantitatively reproduce the measured diffusion of a protein, the simulated result may be corrected using the PSD expression, provided the viscosity of the membrane and the solvent have been determined independently. (Similarly, asking whether lipids may be described by SD-like continuum diffusion requires a periodic correction based on cylinders which span only one leaflet.)

However, a more challenging situation arises if one seeks to compare a simulation result to a more complex experimental system — a phase separated membrane, with regions of different viscosity, or a membrane anchored to a cytoskeleton, or a membrane dense with membrane protein. Here, one requires a simulation approach that

i. is of sufficient scale to encompass the phenomena of interest and so that finite size effects are minimized.
ii. incorporates sufficient chemical accuracy to faithfully represent lipid and protein interactions
iii. includes momentum transport in both membrane and bulk solvent

Item (i) is satisfied by the continuum methods developed by Brown and coworkers discussed above,(57, 63) and other continuum approaches developed to study the dynamics of lipid domains(64) or nonequilibrium dynamics imposed by cortical actin.(65)

Item (ii) is satisfied by use of an all-atom model like Charmm36,(66) which has recently been shown to reproduce 2H NMR quadrupolar spectra for cholesterol rich ternary mixtures.(45) The coarse-grained Martini model, though not quite as accurate at the level of molecular interactions, does reproduce key aspects of lipid phase diagrams.(67) With an explicit representation of the solvent, both methods address item (ii) as well. However, achieving the lengthscales (ca. 100 nm) required to address item (iii) with these models seems intractable for the near future. Such a simulation would contain roughly 100 million particles rendered in all-atom detail, or about 10 million with the coarse-grained Martini model. Note that several authors have reported Martini simulations of protein-dense(68, 69) and actin coupled(70) membranes on ca. 100 nm lengthscales, but in every case these simulations reduce the solvent to a depth of 20 or so nm, and therefore the role of the finite size artifacts discussed above should be carefully considered.(60–62)

Chemical detail in the membrane may be retained at a greatly reduced computational cost with an implicit solvent membrane model like Dry Martini.(71) This fails on item (iii) however, as there is no longer any bulk solvent. To address this, Zgorski and Lyman have recently implemented a fast “Multiparticle Collision” type method for solvent hydrodynamics with Dry Martini, and shown that the approach agrees with the PSD theory of Camley and Brown.(72) The method, “Stochastic Thermostatted Rotation Dynamics (STRD) Martini” uses the Stochastic Rotation Dynamics formulation of MPC to resolve hydrodynamic flows with a resolution of 1-2 nm, but without computation of interparticle distances in the solvent. A STRD Martini simulation of a 100 x 100 nm membrane patch entails molecular dynamics simulation of ca. 400,000 particles, and is therefore readily tractable on modest resources.

In retaining a molecularly resolved membrane, the STRD Martini approach complements methods that treat the membrane as a continuous object. The method of Brown (already mentioned above) tracks a two dimensional representation of the membrane in real space, and propagates hydrodynamic interactions in the Fourier domain, exploiting the simple structure of the quasi2D, Oseen-like tensor. The dynamics of membrane embedded rigid objects are modeled as discrete points. Rather than directly addressing the difficult fluid/object boundary condition, the mobility of the object is obtained numerically from the Green’s functions for fluid flow in the membrane plane, one for each point which resolve the object.(57) As described above, this method has been used to obtain diffusion in solid supported membranes, and in the periodic geometries used in molecular simulation. It is applicable to more complex situations, including bilayer structure and other boundary conditions.(60, 63) The key advantage over particle-based approaches is that much larger membrane areas and timescales are tractable; the main disadvantages are the loss of chemical detail and the need for a Green’s function for the problem of interest, which may be difficult to obtain for more complex membranes.

### Summary

With spatial resolution from 1 to 40 nm and temporal resolution from 1 to 10 microseconds, new experimental techniques are revealing the heterogeneous nature of the plasma membrane on spatiotemporal scales relevant to the dynamics of protein clusters, lipid nanodomains, and within individual actin compartments. At this scale, individual molecular encounters are observed, and it becomes possible to test hypotheses regarding the role of interactions among lipids and proteins in governing membrane organization. However, the observed mobility of membrane proteins depends in complicated and often counterintuitive ways on features such as immobile membrane proteins and regions of differing viscosity. Interpretation of experimental measurements therefore demands modeling approaches which resolve the chemical specificity of membrane interactions, yet remain faithful to the dynamics of lateral diffusion and transport in membranes.

In this effort, the development of model systems intermediate in complexity between cell membranes and simple model systems is essential. Recapitulation of different complicating factors — the actin cytoskeleton, lipidomic complexity, crowding by membrane proteins — in a controlled environment should lead to a more complete understanding of how these features enter the spatiotemporal organization of cell membranes. Comparison of experimental measurements for such systems to simulations designed to incorporate these same features will help to rationalize the measurements, and perhaps support the development of theory which is better suited to the complex environment of the plasma membrane.

## Acknowledgements

E.L. was supported by NIH P20GM104316 and R01 GM120351. JC. E. greatly acknowledges general support by his team and financial support by the MRC (grant number MC_UU_12010/unit programs G0902418 and MC_UU_12025), the Wellcome Trust (grant 104924/14/Z/14 and Strategic Award 091911 (Micron)), MRC/BBSRC/EPSRC (grant MR/K01577X/1), Deutsche Forschungsgemeinschaft (Research unit 1905 “Structure and function of the peroxisomal translocon”), the Wolfson Foundation, the EPA Cephalosporin Fund and the John Fell Fund. C.-L. H. acknowledges support by the Career Development Award, Academia Sinica and by the Ministry of Science and Technology, Taiwan (grant no. 105-2112-M-001-016-MY3)

## References

1. Oldfield, E., and D. Chapman. 1972. Dynamics of Lipids in Membranes: Heterogeneity and the Role of Cholesterol. FEBS letters 23:285–297.

2. Jain, M. K., and H. B. White Iii. 1977. Long-Range Order in Biomembranes. In Advances in Lipid Research. P. Rodolfo, and K. David, editors. Elsevier. 1–60.

3. Simons, K., and E. Ikonen. 1997. Functional rafts in cell membranes. Nature 387:569–572.

4. Feigenson, G. W. 2009. Phase diagrams and lipid domains in multicomponent lipid bilayer mixtures. Biochimica et Biophysica Acta (BBA) - Biomembranes 1788:47–52.

5. Simons, K., and M. J. Gerl. 2010. Revitalizing membrane rafts: new tools and insights. Nature Reviews Molecular Cell Biology 11:688.

6. Sezgin, E., I. Levental, S. Mayor, and C. Eggeling. 2017. The mystery of membrane organization: composition, regulation and roles of lipid rafts. Nat Rev Mol Cell Biol 18:361–374.

7. Manley, S., J. M. Gillette, G. H. Patterson, H. Shroff, H. F. Hess, E. Betzig, and J. Lippincott-Schwartz. 2008. High-density mapping of single-molecule trajectories with photoactivated localization microscopy. Nature Methods 5:155.

8. Kusumi, A., C. Nakada, K. Ritchie, K. Murase, K. Suzuki, H. Murakoshi, R. S. Kasai, J. Kondo, and T. Fujiwara. 2005. Paradigm Shift of the Plasma Membrane Concept from the Two-Dimensional Continuum Fluid to the Partitioned Fluid: High-Speed Single-Molecule Tracking of Membrane Molecules. Annual Review of Biophysics and Biomolecular Structure 34:351–378.

9. Wu, H.-M., Y.-H. Lin, T.-C. Yen, and C.-L. Hsieh. 2016. Nanoscopic substructures of raft-mimetic liquid-ordered membrane domains revealed by high-speed single-particle tracking. Scientific Reports 6:20542.

10. Hsieh, C.-L., S. Spindler, J. Ehrig, and V. Sandoghdar. 2014. Tracking Single Particles on Supported Lipid Membranes: Multimobility Diffusion and Nanoscopic Confinement. The Journal of Physical Chemistry B 118:1545–1554.

11. de Wit, G., J. S. H. Danial, P. Kukura, and M. I. Wallace. 2015. Dynamic label-free imaging of lipid nanodomains. Proceedings of the National Academy of Sciences 112:12299–12303.

12. Huang, Y.-F., G.-Y. Zhuo, C.-Y. Chou, C.-H. Lin, W. Chang, and C.-L. Hsieh. 2017. Coherent Brightfield Microscopy Provides the Spatiotemporal Resolution To Study Early Stage Viral Infection in Live Cells. ACS Nano 11:2575–2585.

13. Schwille, P., U. Haupts, S. Maiti, and W. W. Webb. 1999. Molecular Dynamics in Living Cells Observed by Fluorescence Correlation Spectroscopy with One- and Two-Photon Excitation. Biophysical Journal 77:2251–2265.

14. Wawrezinieck, L., H. Rigneault, D. Marguet, and P.-F. Lenne. 2005. Fluorescence Correlation Spectroscopy Diffusion Laws to Probe the Submicron Cell Membrane Organization. Biophysical Journal 89:4029–4042.

15. Sankaran, J., M. Manna, L. Guo, R. Kraut, and T. Wohland. 2009. Diffusion, Transport, and Cell Membrane Organization Investigated by Imaging Fluorescence Cross-Correlation Spectroscopy. Biophysical Journal 97:2630–2639.

16. Di Rienzo, C., E. Gratton, F. Beltram, and F. Cardarelli. 2013. Fast spatiotemporal correlation spectroscopy to determine protein lateral diffusion laws in live cell membranes. Proceedings of the National Academy of Sciences 110:12307–12312.

17. Regmi, R., P. M. Winkler, V. Flauraud, K. J. E. Borgman, C. Manzo, J. Brugger, H. Rigneault, J. Wenger, and M. F. García-Parajo. 2017. Planar Optical Nanoantennas Resolve Cholesterol-Dependent Nanoscale Heterogeneities in the Plasma Membrane of Living Cells. Nano Letters 17:6295–6302.

18. Eggeling, C., C. Ringemann, R. Medda, G. Schwarzmann, K. Sandhoff, S. Polyakova, V. N. Belov, B. Hein, C. von Middendorff, A. Schonle, and S. W. Hell. 2009. Direct observation of the nanoscale dynamics of membrane lipids in a living cell. Nature 457:1159–1162.

19. Parton, R. G. 2003. Caveolae — from ultrastructure to molecular mechanisms. 4:162.

20. van Zanten, T. S., A. Cambi, M. Koopman, B. Joosten, C. G. Figdor, and M. F. Garcia-Parajo. 2009. Hotspots of GPI-anchored proteins and integrin nanoclusters function as nucleation sites for cell adhesion. Proceedings of the National Academy of Sciences 106:18557–18562.

21. Parasassi, T., E. K. Krasnowska, L. Bagatolli, and E. Gratton. 1998. Laurdan and Prodan as Polarity-Sensitive Fluorescent Membrane Probes. Journal of Fluorescence 8:365–373.

22. Sezgin, E., and P. Schwille. 2012. Model membrane platforms to study protein-membrane interactions. Molecular Membrane Biology 29:144–154.

23. Gerl, M. J., J. L. Sampaio, S. Urban, L. Kalvodova, J.-M. Verbavatz, B. Binnington, D. Lindemann, C. A. Lingwood, A. Shevchenko, C. Schroeder, and K. Simons. 2012. Quantitative analysis of the lipidomes of the influenza virus envelope and MDCK cell apical membrane. The Journal of Cell Biology 196:213–221.

24. Sampaio, J. L., M. J. Gerl, C. Klose, C. S. Ejsing, H. Beug, K. Simons, and A. Shevchenko. 2011. Membrane lipidome of an epithelial cell line. Proceedings of the National Academy of Sciences 108:1903–1907.

25. Tulodziecka, K., B. B. Diaz-Rohrer, M. M. Farley, R. B. Chan, G. Di Paolo, K. R. Levental, M. N. Waxham, and I. Levental. 2016. Remodeling of the postsynaptic plasma membrane during neural development. Molecular Biology of the Cell 27:3480–3489.

26. Sezgin, E., T. Gutmann, T. Buhl, R. Dirkx, M. Grzybek, Ü. Coskun, M. Solimena, K. Simons, I. Levental, and P. Schwille. 2015. Adaptive Lipid Packing and Bioactivity in Membrane Domains. PLoS ONE 10:e0123930.

27. Spendier, K., K. A. Lidke, D. S. Lidke, and J. L. Thomas. 2012. Single-particle tracking of immunoglobulin E receptors (FcεRI) in micron-sized clusters and receptor patches. FEBS Letters 586:416–421.

28. Fujiwara, T., K. Ritchie, H. Murakoshi, K. Jacobson, and A. Kusumi. 2002. Phospholipids undergo hop diffusion in compartmentalized cell membrane. The Journal of Cell Biology 157:1071.

29. Andrade, D. M., M. P. Clausen, J. Keller, V. Mueller, C. Wu, J. E. Bear, S. W. Hell, B. C. Lagerholm, and C. Eggeling. 2015. Cortical actin networks induce spatio-temporal confinement of phospholipids in the plasma membrane – a minimally invasive investigation by STED-FCS. Scientific Reports 5:11454.

30. Sadegh, S., J. L. Higgins, P. C. Mannion, M. M. Tamkun, and D. Krapf. 2017. Plasma Membrane is Compartmentalized by a Self-Similar Cortical Actin Meshwork. Physical Review X 7:011031.

31. Machta, Benjamin B., S. Papanikolaou, James P. Sethna, and Sarah L. Veatch. 2011. Minimal Model of Plasma Membrane Heterogeneity Requires Coupling Cortical Actin to Criticality. Biophysical Journal 100:1668–1677.

32. Honigmann, A., S. Sadeghi, J. Keller, S. W. Hell, C. Eggeling, R. Vink, and R. Schekman. 2014. A lipid bound actin meshwork organizes liquid phase separation in model membranes. eLife 3.

33. Heinemann, F., Sven K. Vogel, and P. Schwille. 2013. Lateral Membrane Diffusion Modulated by a Minimal Actin Cortex. Biophysical Journal 104:1465–1475.

34. Manzo, C., T. S. van Zanten, and M. F. Garcia-Parajo. 2011. Nanoscale Fluorescence Correlation Spectroscopy on Intact Living Cell Membranes with NSOM Probes. Biophysical Journal 100:L8–L10.

35. Winkler, P. M., R. Regmi, V. Flauraud, J. Brugger, H. Rigneault, J. Wenger, and M. F. García-Parajo. 2017. Transient Nanoscopic Phase Separation in Biological Lipid Membranes Resolved by Planar Plasmonic Antennas. ACS Nano 11:7241–7250.

36. Ringemann, C., B. Harke, C. v. Middendorff, R. Medda, A. Honigmann, R. Wagner, M. Leutenegger, A. Schönle, S. W. Hell, and C. Eggeling. 2009. Exploring single-molecule dynamics with fluorescence nanoscopy. New Journal of Physics 11:103054.

37. Eggeling, C. 2015. Super-resolution optical microscopy of lipid plasma membrane dynamics. Essays In Biochemistry 57:69.

38. Vicidomini, G., H. Ta, A. Honigmann, V. Mueller, M. P. Clausen, D. Waithe, S. Galiani, E. Sezgin, A. Diaspro, S. W. Hell, and C. Eggeling. 2015. STED-FLCS: An Advanced Tool to Reveal Spatiotemporal Heterogeneity of Molecular Membrane Dynamics. Nano Letters 15:5912–5918.

39. Honigmann, A., V. Mueller, H. Ta, A. Schoenle, E. Sezgin, S. W. Hell, and C. Eggeling. 2014. Scanning STED-FCS reveals spatiotemporal heterogeneity of lipid interaction in the plasma membrane of living cells. Nat Commun 5:1–11.

40. Mueller, V., C. Ringemann, A. Honigmann, G. Schwarzmann, R. Medda, M. Leutenegger, S. Polyakova, V. N. Belov, S. W. Hell, and C. Eggeling. 2011. STED Nanoscopy Reveals Molecular Details of Cholesterol- and Cytoskeleton-Modulated Lipid Interactions in Living Cells. Biophysical Journal 101:1651–1660.

41. Schneider, F., D. Waithe, M. P. Clausen, S. Galiani, T. Koller, G. Ozhan, C. Eggeling, and E. Sezgin. 2017. Diffusion of lipids and GPI-anchored proteins in actin-free plasma membrane vesicles measured by STED-FCS. Molecular Biology of the Cell 28:1507–1518.

42. Honigmann, A., V. Mueller, S. W. Hell, and C. Eggeling. 2013. STED microscopy detects and quantifies liquid phase separation in lipid membranes using a new far-red emitting fluorescent phosphoglycerolipid analogue. Faraday Discussions 161:77–89.

43. Lindfors, K., T. Kalkbrenner, P. Stoller, and V. Sandoghdar. 2004. Detection and Spectroscopy of Gold Nanoparticles Using Supercontinuum White Light Confocal Microscopy. Physical Review Letters 93:037401.

44. Kukura, P., H. Ewers, C. Muller, A. Renn, A. Helenius, and V. Sandoghdar. 2009. High-speed nanoscopic tracking of the position and orientation of a single virus. Nat Meth 6:923–927.

45. Sodt, A. J., M. L. Sandar, K. Gawrisch, R. W. Pastor, and E. Lyman. 2014. The Molecular Structure of the Liquid-Ordered Phase of Lipid Bilayers. Journal of the American Chemical Society 136:725–732.

46. Sodt, A. J., R. W. Pastor, and E. Lyman. 2015. Hexagonal Substructure and Hydrogen Bonding in Liquid Ordered Phases of Palmitoyl Sphingomyelin. Biophys. J. 109:948–955.

47. Hsieh, C.-L. 2018. Label-free, ultrasensitive, ultrahigh-speed scattering-based interferometric imaging. Optics Communications, in press.

48. Cheng, C.-Y., and C.-L. Hsieh. 2017. Background Estimation and Correction for High-Precision Localization Microscopy. ACS Photonics 4:1730–1739.

49. Clausen, M. P., and B. C. Lagerholm. 2011. The Probe Rules in Single Particle Tracking. Current Protein & Peptide Science 12:699–713.

50. Cole, D., G. Young, A. Weigel, A. Sebesta, and P. Kukura. 2017. Label-Free Single-Molecule Imaging with Numerical-Aperture-Shaped Interferometric Scattering Microscopy. ACS Photonics 4:211–216.

51. Liebel, M., J. T. Hugall, and N. F. van Hulst. 2017. Ultrasensitive Label-Free Nanosensing and High-Speed Tracking of Single Proteins. Nano Letters 17:1277–1281.

52. Hedde, P. N., R. M. Dörlich, R. Blomley, D. Gradl, E. Oppong, A. C. B. Cato, and G. U. Nienhaus. 2013. Stimulated emission depletion-based raster image correlation spectroscopy reveals biomolecular dynamics in live cells. Nature Communications 4:2093.

53. Heine, J., M. Reuss, B. Harke, E. D’Este, S. J. Sahl, and S. W. Hell. 2017. Adaptive-illumination STED nanoscopy. Proceedings of the National Academy of Sciences 114:9797.

54. Saffman, P. G., and M. Delbrück. 1975. Brownian motion in biological membranes. Proceedings of the National Academy of Sciences 72:3111–3113.

55. Petrov, E. P., and P. Schwille. 2008. Translational Diffusion in Lipid Membranes beyond the Saffman-Delbrück Approximation. Biophysical Journal 94:L41–L43.

56. Knight, J. D., M. G. Lerner, J. G. Marcano-Velázquez, R. W. Pastor, and Joseph J. Falke. Single Molecule Diffusion of Membrane-Bound Proteins: Window into Lipid Contacts and Bilayer Dynamics. Biophysical Journal 99:2879–2887.

57. Camley, B. A., and F. L. H. Brown. 2013. Diffusion of complex objects embedded in free and supported lipid bilayer membranes: role of shape anisotropy and leaflet structure. Soft Matter 9:4767–4779.

58. Oppenheimer, N., and H. Diamant. 2009. Correlated Diffusion of Membrane Proteins and Their Effect on Membrane Viscosity. Biophysical Journal 96:3041–3049.

59. Oppenheimer, N., and H. Diamant. 2011. In-Plane Dynamics of Membranes with Immobile Inclusions. Physical Review Letters 107:258102.

60. Camley, B. A., M. G. Lerner, R. W. Pastor, and F. L. H. Brown. 2015. Strong influence of periodic boundary conditions on lateral diffusion in lipid bilayer membranes. The Journal of Chemical Physics 143:243113.

61. Venable, R. M., H. I. Ingólfsson, M. G. Lerner, B. S. Perrin, B. A. Camley, S. J. Marrink, F. L. H. Brown, and R. W. Pastor. 2016. Lipid and Peptide Diffusion in Bilayers: The Saffman–Delbrück Model and Periodic Boundary Conditions. The Journal of Physical Chemistry B.

62. Vögele, M., and G. Hummer. 2016. Divergent Diffusion Coefficients in Simulations of Fluids and Lipid Membranes. The Journal of Physical Chemistry B 120:8722–8732.

63. Noruzifar, E., B. A. Camley, and F. L. H. Brown. 2014. Calculating hydrodynamic interactions for membrane-embedded objects. The Journal of Chemical Physics 141:124711.

64. Fan, J., T. Han, and M. Haataja. 2010. Hydrodynamic effects on spinodal decomposition kinetics in planar lipid bilayer membranes. The Journal of Chemical Physics 133:235101.

65. Goswami, D., K. Gowrishankar, S. Bilgrami, S. Ghosh, R. Raghupathy, R. Chadda, R. Vishwakarma, M. Rao, and S. Mayor. 2008. Nanoclusters of GPI-Anchored Proteins Are Formed by Cortical Actin-Driven Activity. Cell 135:1085–1097.

66. Klauda, J. B., R. M. Venable, J. A. Freites, J. W. O’Connor, D. J. Tobias, C. Mondragon-Ramirez, I. Vorobyov, A. D. MacKerell, and R. W. Pastor. 2010. Update of the CHARMM All-Atom Additive Force Field for Lipids: Validation on Six Lipid Types. The Journal of Physical Chemistry B 114:7830–7843.

67. Ingólfsson, H. I., M. N. Melo, F. J. van Eerden, C. Arnarez, C. A. Lopez, T. A. Wassenaar, X. Periole, A. H. de Vries, D. P. Tieleman, and S. J. Marrink. 2014. Lipid Organization of the Plasma Membrane. Journal of the American Chemical Society 136:14554–14559.

68. Arnarez, C., S. J. Marrink, and X. Periole. 2016. Molecular mechanism of cardiolipin-mediated assembly of respiratory chain supercomplexes. Chemical Science 7:4435–4443.

69. Javanainen, M., H. Martinez-Seara, R. Metzler, and I. Vattulainen. 2017. Diffusion of Integral Membrane Proteins in Protein-Rich Membranes. The Journal of Physical Chemistry Letters 8:4308–4313.

70. Koldsø, H., T. Reddy, P. W. Fowler, A. L. Duncan, and M. S. P. Sansom. 2016. Membrane Compartmentalization Reducing the Mobility of Lipids and Proteins within a Model Plasma Membrane. The Journal of Physical Chemistry B 120:8873–8881.

71. Arnarez, C., J. J. Uusitalo, M. F. Masman, H. I. Ingólfsson, D. H. de Jong, M. N. Melo, X. Periole, A. H. de Vries, and S. J. Marrink. 2015. Dry Martini, a Coarse-Grained Force Field for Lipid Membrane Simulations with Implicit Solvent. Journal of Chemical Theory and Computation 11:260–275.

72. Zgorski, A., and E. Lyman. 2016. Toward Hydrodynamics with Solvent Free Lipid Models: STRD Martini. Biophysical Journal 111:2689–2697.

